# Open-3DSIM: an Open-source three-dimensional structured illumination microscopy reconstruction platform

**DOI:** 10.1101/2022.12.16.520543

**Authors:** Ruijie Cao, Yaning Li, Xin Chen, Xichuan Ge, Meiqi Li, Meiling Guan, Yiwei Hou, Yunzhe Fu, Shan Jiang, Baoxiang Gao, Peng Xi

## Abstract

With optical section and defocus removal effect, three-dimensional structured illumination microscopy (3DSIM) can get a whole sight of intracellular organelle. Here, Open-3DSIM is reported as an open-source reconstruction platform with double improvement on lateral and axial resolution. MATLAB code, ImageJ version and Exe application are provided for biologists and engineers to maximize its user-friendliness and prompt its further development. Through adaptive parameter estimation and spectrum filter optimization, we demonstrate its superior performance of artifact suppression and defocus elimination over other algorithms on various specimens, under gradient signal-to-noise levels. Moreover, with the capacity to extract the dipole orientation, Open-3DSIM paves a new avenue for interpreting the subcellular structures in six dimensions (*xyzθλt*).

## Introduction

Structured illumination microscopy is the *de facto* most universally implemented super-resolution microscopy in visualizing intracellular organelle and cytoskeletal interactions, due to its low photo-toxicity, fast imaging speed, long time-lapse, and high compatibility with existing fluorescent labelling^1-3^. With the flourishing of SIM developments, a variety of open-source and high-fidelity reconstruction algorithms have been developed, such as OpenSIM^4^, fairSIM^5^, SIMtoolbox^6^, HiFi-SIM^7^, etc. Of them, OpenSIM is a classical two-dimensional structure illumination microscopy (2DSIM) reconstruction algorithm based on MATLAB, fairSIM provides a reconstruction of 2DSIM and single-layer three-dimensional SIM (3DSIM) based on ImageJ, SIMToolbox includes optical section SIM and 2DSIM in MATLAB, and HiFi-SIM optimizes reconstruction performance of 2DSIM and single-layer 3DSIM by engineering the effective point spread function in MATLAB. As SIM requires frequency-domain operations to minimize parameter estimation errors and artifacts of reconstructed images, the customized open-source tools with empirical parameter adjustments often outperform the automated commercial software.

The development of the open-source SIM reconstruction software also boosts custom-built SIM hardware platforms, such as SLM-SIM^8^, DMD-SIM^9^, galvanometer-SIM^10^, Hessian-SIM^11^, Multi-SIM^12^, etc. However, most existing SIM software can only reconstruct 2D or single-layer 3DSIM data by shifting the spectrum in 2D frequency domain such as fairSIM and HiFi-SIM^13, 14^. The existing multi-layer SIM algorithms are either provided as closed-source commercial 3DSIM systems such as GE Deltavision OMX SR (OMX)^15^, or as purpose-written tools with time- consuming optimization and non-universality by research groups such as noise- control SIM reconstruction (SIMnoise)^16^. Lack of open-source and user-friendly software hinders the users to access and use 3DSIM, thus it makes the development of the corresponding hardware far behind its sibling 2DSIM.

To answer this call, here we report Open-3DSIM, which can provide superior and robust multi-layer 3DSIM reconstruction. We compare its performance with OMX and SIMnoise on various specimens and gradient signal-to-noise (SNR) levels. Results show that through optimization on parameter estimation and spectral filter, Open-3DSIM outperforms other algorithms in respect of high fidelity, low artifact, and defocus-removal. Furthermore, Open-3DSIM can extract the inherent dipole orientation information embedded in the existing 3D-SIM data. It enables the multi- layer, multi-color, polarization, time-lapse super-resolution SIM reconstruction, unlocking the full potential of SIM in six dimensions (*xyzθλt*). To make Open-3DSIM a user-friendly tool, we prepared the ImageJ version and Exe application with detailed guidelines and test datasets to make it friend to biological users. And, we also provide MATLAB codes for algorithm users to boost future developments of 3DSIM algorithms such as frame-reduction, machine learning, or post-processing based on regularity/prior knowledge.

### Comparison of single-layer SIM and multi-layer SIM

Although 2D-SIM has achieved double improvement on *xoy* plane, but the axial resolution is not improved. This results in severe defocus background and artifacts as shown in **Fig. 1(a)** and **(b)**. Although HiFi-SIM achieves good effect of artifact removal and fidelity, it is still restricted in single-layer. When it comes to thick samples with serious defocused backgrounds, the reconstructed results of HiFi-SIM have more defocused backgrounds and defocused artifacts with no improvement in *xoz* resolution. By comparison, Open-3DSIM can better remove the defocus or artifacts, improve axial resolution by fill the leaky-cone of optical transfer function (OTF). **Fig. 1(c)** shows the 3D-SIM imaging of actin filaments in U2OS. It can be seen that Open-3DSIM can improve 3D resolution compared to wide-field image. And, Open-3DSIM interpreting whole actin structure can greatly eliminate the influence of defocus, background and artifacts compared with single-layer reconstruction. We also introduce polarization dimension on *xoy* plane to realize dipole orientation imaging in Open-3DSIM^18, 19^. It’s noteworthy that with intensity calibration, the dipole orientation will be more accurately resolved by Open-3DSIM. The parallelism between dipole orientation and actin filament direction validates the correctness of polarization information.

**Fig. 1.**
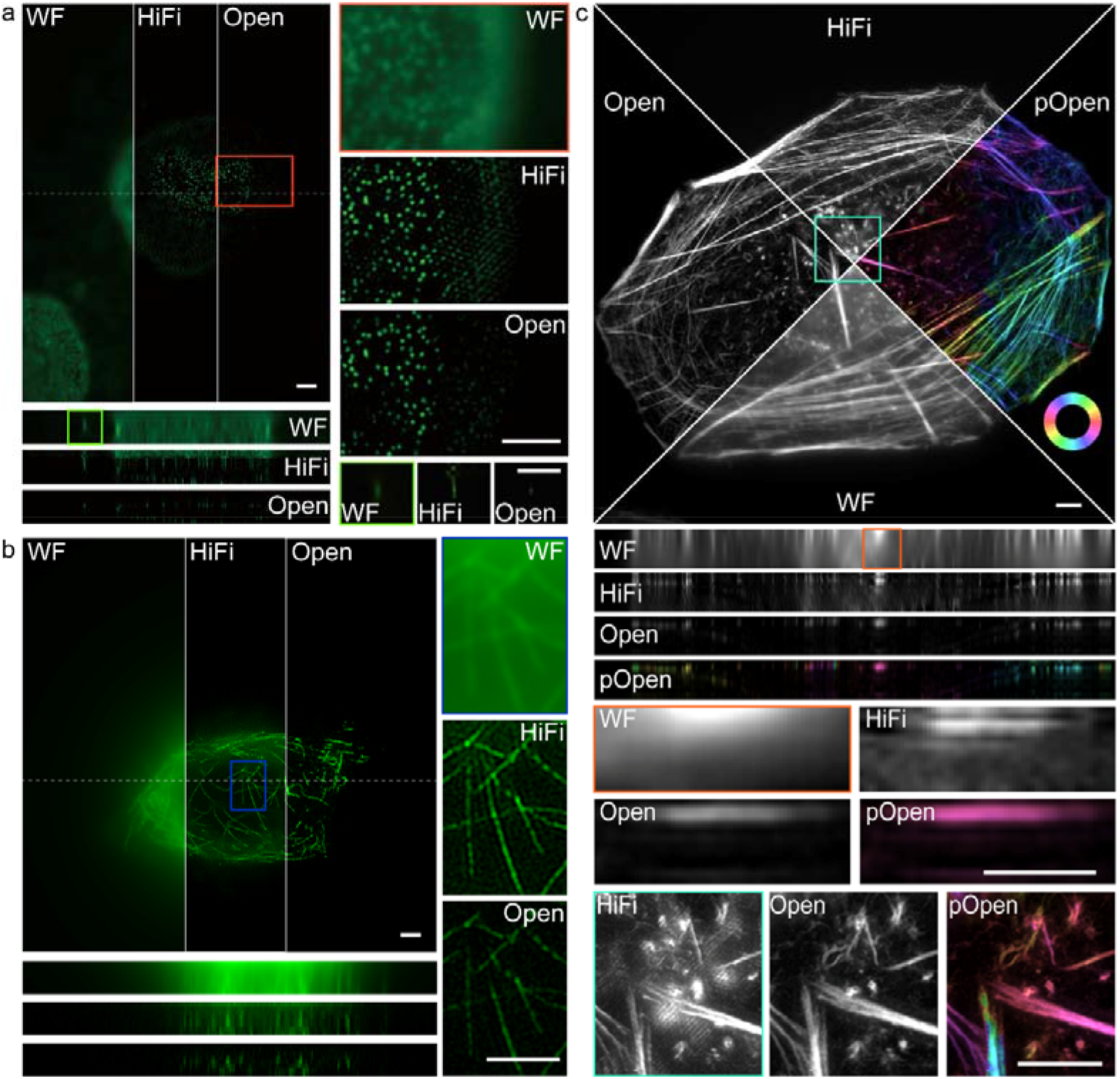
Comparison between single-layer SIM and multi-layer SIM. The comparison of the single-layer algorithm (HiFi-SIM), the multi-layer algorithm (Open-3DSIM), widefield (WF) on **a**, nuclear pore complex, **b**, actin filament in U2OS osteosarcoma cells from fairSIM, and **c**, actin filaments in U2OS, including polarized Open-3DSIM image (pOpen). Scale bar: 2 μm.

### Adaptive parameter estimation and spectrum optimization

We use cross-correlation method^5, 7^ to estimate the frequency, angle, phase, and modulation depth of the structured illumination patterns. To obtain a correct frequency parameter which is the first step of parameter estimation, we use both +1^st^ and +2^nd^ order frequencies. Although traditionally estimating the peak of +2^th^ frequency is more accurate than +1^st^, +1^st^ frequency’s peak carries a higher contrast than +2^nd^ frequency in low SNR as shown in **Fig. 2(a)**. Therefore, we set up a criterion to determine whether estimating through +2^nd^ frequency’s peak is reliable. If not, we will use the +1^st^ spectrum to estimate instead to guarantee the validity of frequency estimation. Then the corresponding parameter such as phase, angle can be accurately resolved. This method can greatly improve the correctness of parameter estimation when reconstructing images under low SNR, thus reducing various artifacts caused by parameter estimation errors^10, 17^.

**Fig. 2.**
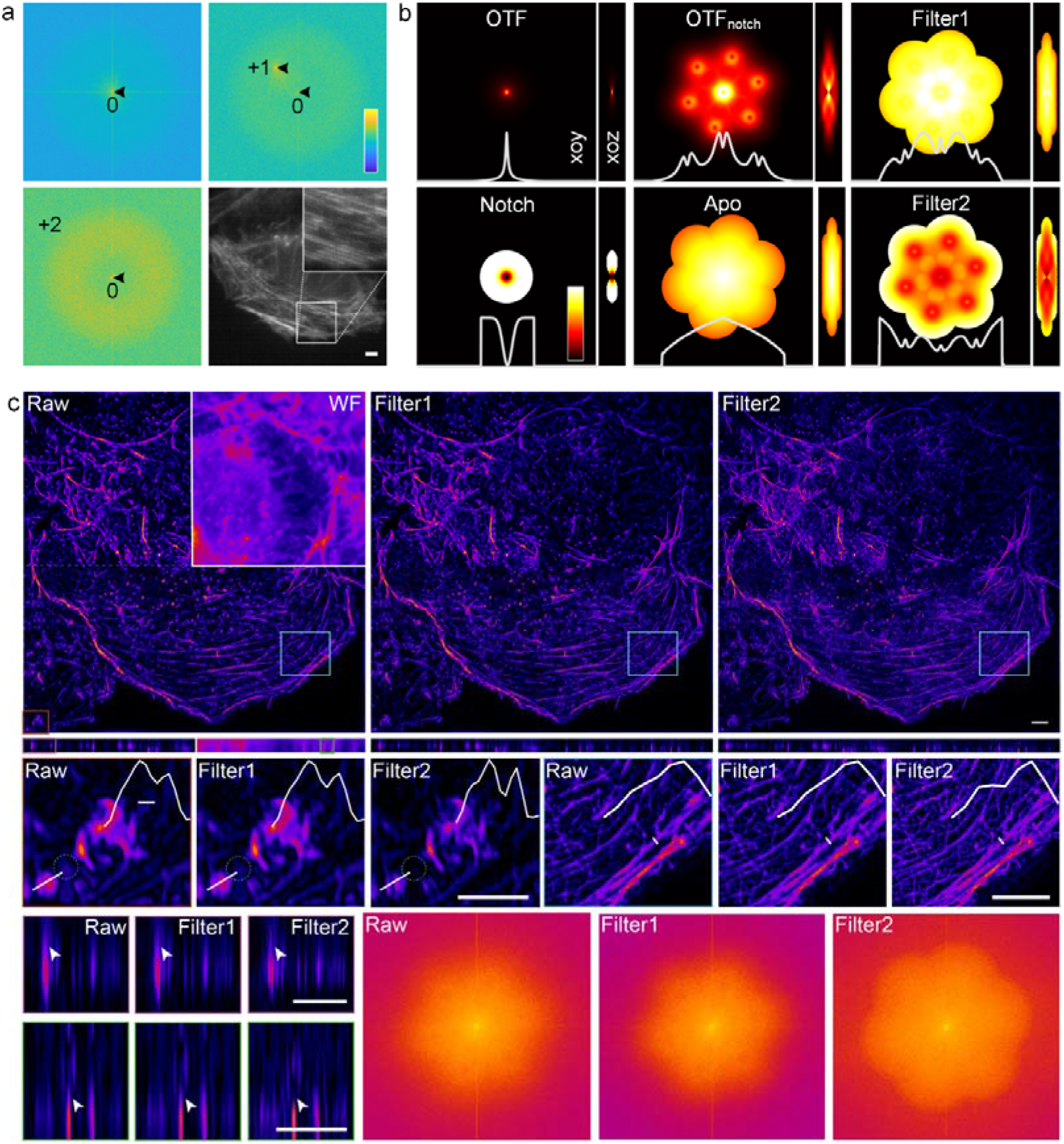
Adaptive estimation and spectrum optimization. **a**, Separated frequency domain and image under low SNR, demonstrating the 1^st^ frequency peak is higher than the 2^nd^ and is easier to recognize. Top left, top right and bottom left are the separated 0^th^, 1^st^, and 2^nd^ frequency domain. Bottom right is the raw image of actin filaments in U2OS under extremely low SNR, but the +1^st^ pattern (visible because of relative low frequency) is easy to recognize. **b**, Two-step filter designed by OTF, Notch, OTF_notch,_ and Apodization to optimize the frequency spectrum of 3DSIM, achieving the effect of reducing artifacts and improving resolution. The white lines below represent their corresponding profiles along the estimated frequency.

Next, according to the estimated frequency vector in *xoy* and *yoz* plane, we designed a two-step filter in frequency domain based on notch function (Notch), apodization function (Apo), optical conversion function (OTF), and OTF_notch_ to suppress spectrum’s peak, reduce the noise and compensate high-frequency information after OTF blurring, as shown in **Fig. 2(b)**. We design two filters of Filter1 and Filter2 to suppress the artifacts of reconstructed images and further improve the 3D resolution respectively. We demonstrate that Filter1 can greatly reduce the artifact and Filter2 can improve the 3D resolution of reconstruction images in **Fig. 2(c)**.

### Open-3DSIM demonstrates superior reconstruction

To further test the performance of Open-3DSIM, we compare the reconstruction results of Open-3DSIM, OMX, and SIMnoise under gradient SNR for actin filaments in U2OS in **Fig.3(a)**. Here, Open-3DSIM outperforms OMX and SIMnoise in different levels of SNR with less artifacts and background. And the mean absolute error (MAE) and SNR of Open-3DSIM are almost the same as those of other algorithms with a higher SNR in **Fig. 3(b)**, proving the excellent performance of Open-3DSIM under low SNR. Although SIMnoise has optimized the reconstruction under low SNR, Open-3DSIM shows better edge information retention and denoising effect compared with the state-of-art result of SIMnoise in **Fig. 3(c)** with less reconstruction time. Similarly, compared with OMX, Open-3DSIM reconstruction algorithm has excellent ability to remove artifacts and retain weak information in **Fig. 3(d)** under low SNR.

**Fig. 3.**
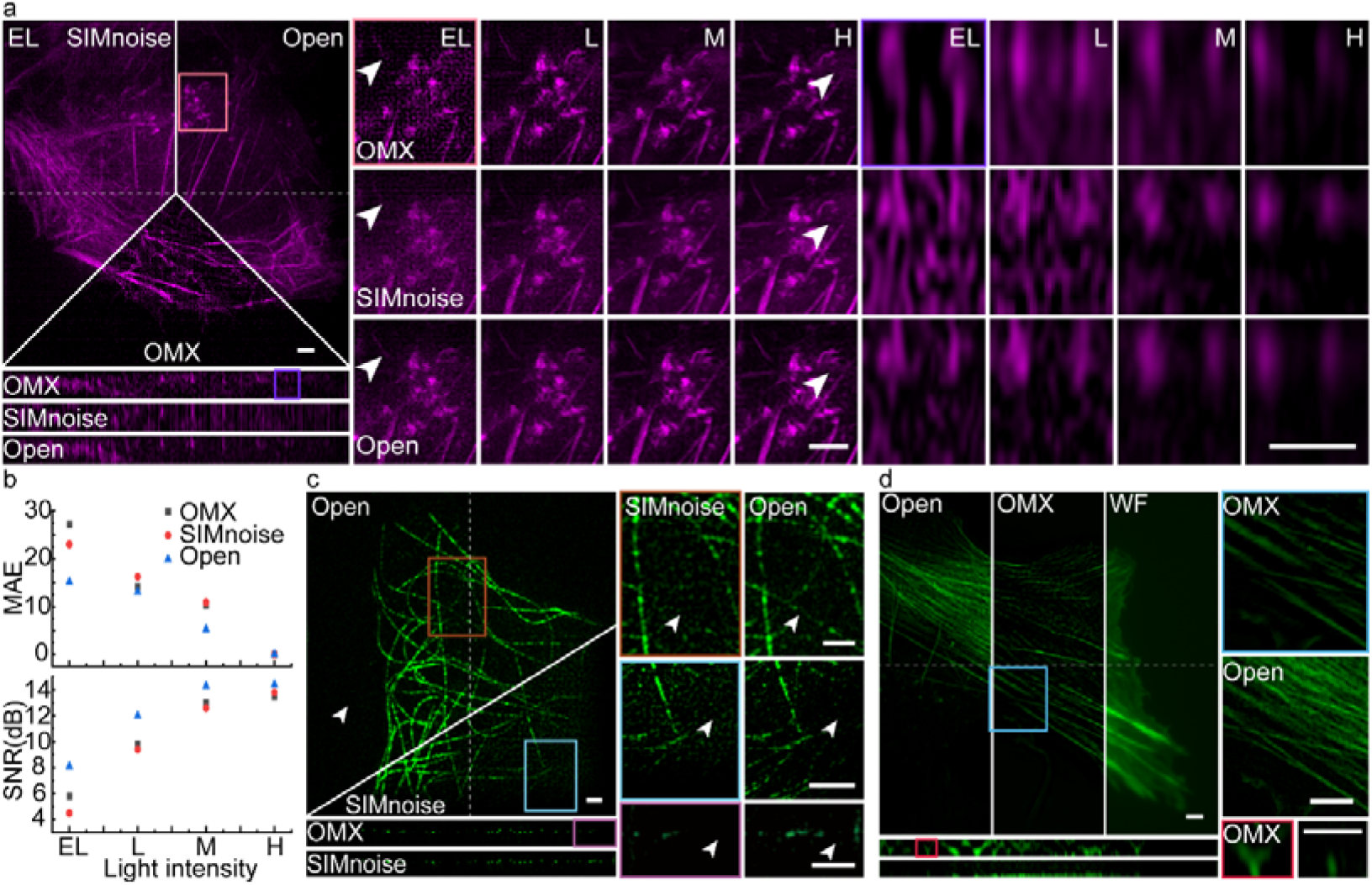
Superior reconstruction performance of Open-3DSIM. **a**, Reconstruction for actin filaments under different SNRs of extremely low (EL, intensity of 2%, exposure of 5ms), low (L, 5%, 5ms), moderate (M, 10%, 20ms), and high (H, 10%, 50ms) with the comparison among SIMnoise, OMX- SIM, and Open-3DSIM. **b**, MAE under different SNRs and algorithms, taking the corresponding high SNR results as ground truth images, with the relative SNR of the reconstruction results under different SNRs and algorithms to demonstrate the algorithm’s ability to suppress defocused background. **c**, State-of-art reconstruction results of tubulin structure from SIMnoise under lowest SNR with the corresponding comparison. **d**, Reconstruction for actin filaments from OMX under low SNR (5%, 20ms) with the corresponding comparison. Scale bar: 2μm.

## Discussion and conclusion

With the effort of the 2DSIM open-source community, the development of a variety of 2DSIM in both hardware and software have been emerging in recent years. Yet, the lack of 3DSIM reconstruction software creates huge obstacle for its technical developments. Especially, due to the tragic discontinuity of GE DeltaVision OMX system, the users need the effort from open-source community to continue such support. With excellent performance on high fidelity, low artifact and defocus- removal on different samples under gradient SNR levels, Open-3DSIM can truly unleash the full potential of the SIM in both lateral and axial super-resolution, through multi-color, multi-layer, time-lapse, and dipole orientation imaging. We expect Open- 3DSIM to be the open-source standard for the 3DSIM multi-dimensional reconstruction, and we believe that the open-source effort will significantly boost the development of custom-built 3DSIM systems, making a great difference to the community.

## Methods

### Sample preparation

Human osteosarcoma U2OS cell lines (HTB-96, ATCC, USA) were cultured in Dulbecco’s Modified Eagle’s medium (DMEM, GIBCO, USA) containing 10% heat-inactivated fetal bovine serum (FBS, GIBCO, USA) and 100□U/ml penicillin and 100□µg/ml streptomycin solution (PS, GIBCO, USA) at 37□°C in an incubator with 95% humidity and 5% CO_2_. Fixed cells were plated on #1.5H cover glasses (CG15XH, Thorlabs, USA) before they were grown to a suitable density (24□hours) and fixed with 4% formaldehyde (R37814, Invitrogen, USA) for 15□min. Then, the cells were washed three times with PBS to remove the formaldehyde. Alexa Flour 488 Phalloidin (A12379, Invitrogen, USA) was used to stain the actin filaments for one hour at room temperature. Then, the coverslip was sealed on the slide with the prolong antifade mountant (P36941, Invitrogen, USA).

The COS7 cells were cultured in Dulbecco’s Modified Eagle’s medium (DMEM, GIBCO, USA) containing 10% (V/V) fetal bovine serum (FBS, GIBCO, USA), and 100□U/ml penicillin and 100□µg/ml streptomycin solution (PS, GIBCO, USA) at 37□in an incubator with 95% humidity atmosphere and 5% CO_2_. The selected cell lines were seeded on #1.5H cover glasses (CG15XH, Thorlabs, USA) before they were grown to a suitable density (24□hours) via incubation in a humid atmosphere containing 5% (V) CO2 at 37□°C. The cells were stained in DMEM containing 500nM Mito tracker™ Orange (M7510, ThermoFisher, USA) and 0.5% dimethyl sulfoxide for 20min in a CO_2_ incubator. The cells were washed with PBS and fixed in 4% formaldehyde (R37814, Invitrogen, USA) for 15□min at room temperature. The coverslip was sealed onto the cavity of the slide. The coverslip was sealed on the slide with the prolong antifade mountant (P36941, Invitrogen, USA). For COS7 living cells imaging, the COS7 cells were seeded on µ-Slide 8 Wellhigh (80806, ibidi, German). The COS7 cells were stained in DMEM containing 500nM IMMBright660 and 0.5% dimethyl sulfoxide for 20min in a CO_2_ incubator. Then, the cells were kept in DMEM for live-cells imaging without washing.

The mouse kidney sections to reconstruct the actin filaments were purchased from ThermoFisher. Actin filaments were labeled by Alexa Fluor 568 using Prolong Diamond to embed.

The COS7 cells to reconstruct the nuclear pore complex were purchased from GATTA Quant. Anti-Nup was labeled by Alexa Fluor 555 using prolong diamond to embed.

The BPAEC cells to reconstruct the multi-color sample were purchased from ThermoFisher. Mitochondria were labeled with red-fluorescent MitoTracker Red CMXRos, F-actin was stained using green-fluorescent Alexa Fluor 488 phalloidin, and blue-fluorescent DAPI was used to label the nuclei using Prolong Diamond to embed.

### Data acquisition

We obtain data mainly based on the commercial OMX-SIM system (DeltaVision OMX SR, GE, USA) using a oil immersion objective (Olympus, Japan, ×60 1.4 NA). 3D-SIM sequences were performed with pixel size of 80nm and 125nm in *xoy* and *xoz* plane (5 phases, 3 angles, and 15 raw images per plane). Samples in **Fig. 1(b), Fig. 2(c)** are obtained from open-source data in fairSIM: http://www.fairsim.org/. Samples in **Fig. 3(c)** is obtained from open-source data in SIMnoise: https://data.4tu.nl/articles/_/12942932.

### Raw image format

Open-3DSIM supports the input document format of *.dv/*.tif/*.tiff. The picture sequence is: phase-depth-channel-time-angle. We suggest using samples of five or more layers for reconstruction to obtain optimal results.

### Image processing

The frequency domain and SNR of images are generated by ImageJ. HiFi-SIM, SIMnoise and commercial OMX-SR algorithm are used for comparison. The reconstructions of Open3DSIM were processed with MATLAB (2018a, 2021a and 2021b) program. Figures were plotted with Origin, Visio and Adobe Illustrator.

### Calibration of the illumination nonuniformity

Fluorescent beads (Thermo Flsher Scientific) with 200 nm in diameter was prepared in a fixed slide. The beads were imaged by OMX for three angles with five phases in each angle. By averaging images of five phases, WF images of each angle can be obtained. To correct the light intensity at each point, the beads should be thick on the focal plane. Then interpolation is used to compensate the gap of beads to form a light intensity map of *calib*_*ang*1_, *calib*_*ang*2_ and *calib*_*ang*3_

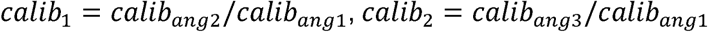

### Calibration of MAE and SNR

MAE and SNR in **Fig.2(b)** is used to evaluate the reconstruction of Open-3DSIM, SIMnoise and OMX-SIM. After image calibration to compensate for vibrating during shooting, the corresponding results of high SNR under different algorithms are used as ground-truth *y*(*i*), so the images of low SNR *y*′(*i*) are calculated using MAE:

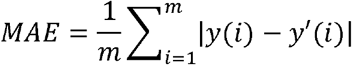

Where *i* denotes the *i*-th pixel of whole *m* pixels. And SNR is used to evaluate the ability to remove defocused background of different algorithm. We choose the part with signal and the part without signal to solve its mean value and variance, and use the following formula to solve SNR (dB):

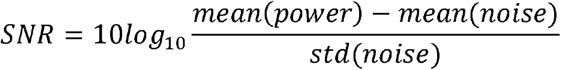

## Supplementary software and data

Additional software has been uploaded to GitHub (https://github.com/Cao-ruijie/Open3DSIM) with detailed user guide and test data. And the additional data has been uploaded to Fishare with dataset, parameters, and comparison. (https://figshare.com/articles/dataset/Open_3DSIM_DATA/21731315).

## Author Contributions

R.C. and P.X. conceived the idea of Open-3DSIM. P.X. supervised the research. R.C., P.X., Y.L., X.C., and M. G. deduce the theory of Open-3DSIM. R.C. and M.L. designed and conduct the majority of the experiments. M.L., X.G, B.G., S. J. and Y.F. prepared the sample. R.C., P.X., Y.L., X.C., Y.H., M.L., and M.G. discussed the theory of Open-3DSIM. R.C. and X.C. discussed the optimization of parameter estimating, and R.C. and Y.L. conducted data spectrum optimization. R.C write the MATLAB and ImageJ version of Open-3DSIM. R.C., Y.L, M.L., Y.H., and P.X. wrote the paper with inputs from all authors.

## Acknowledgments

This work was supported by the National Key R&D Program of China (2022YFC3401100), and National Natural Science Foundation of China (62025501, 31971376, 92150301),. We thank National Center for Protein Sciences at Peking University in Beijing, China, for assistance with GE DeltaVision OMX super-resolution imaging.

## Competing interests

The authors declare no competing financial interests.

## Notes

https://github.com/Cao-ruijie/Open3DSIM

https://figshare.com/articles/dataset/Open_3DSIM_DATA/21731315

### Competing Interest Statement

The authors have declared no competing interest.

## Reference

1. Gustafsson, Mats G L. Surpassing the lateral resolution limit by a factor of two using structured illumination microscopy. Journal of microscopy. Journal of microscopy 198, 82–87 (2000).

2. Rust M J, Bates M, Zhuang X. Sub-diffraction-limit imaging by stochastic optical reconstruction microscopy (STORM). Nature methods 3, 793–796 (2006).

3. Klar T A, Jakobs S, Dyba M, et al. Fluorescence microscopy with diffraction resolution barrier broken by stimulated emission. Proceedings of the National Academy of Sciences 97, 8206–8210 (2000).

4. Lal A, Shan C, Xi P. Structured illumination microscopy image reconstruction algorithm. IEEE Journal of Selected Topics in Quantum Electronics 22, 50–63 (2016).

5. Müller, Marcel, et al. Open-source image reconstruction of super-resolution structured illumination microscopy data in ImageJ. Nature communications 7, 1–6 (2016).

6. Křížek, Pavel, et al. SIMToolbox: a MATLAB toolbox for structured illumination fluorescence microscopy. Bioinformatics 32, 318–320 (2016).

7. Wen, G. et al. High-fidelity structured illumination microscopy by point-spread-function engineering. Light: Science & Applications 10, 1–12 (2021).

8. Xu, L. et al. Structured illumination microscopy based on asymmetric three-beam interference. Journal of Innovative Optical Health Sciences 14, 2050027 (2021).

9. Li, M. et al. Structured illumination microscopy using digital micro-mirror device and coherent light source. Applied Physics Letters 116, 233702 (2020).

10. Liu, W. et al. Three-dimensional super-resolution imaging of live whole cells using galvanometer-based structured illumination microscopy. Optics express 27, 7237–7248 (2019).

11. Huang, X. et al. Fast, long-term, super-resolution imaging with Hessian structured illumination microscopy. Nature biotechnology 36, 451–459 (2018).

12. Aguilar-Martinez, E. et al. Screen for multi-SUMO–binding proteins reveals a multi-SIM–binding mechanism for recruitment of the transcriptional regulator ZMYM2 to chromatin. Proceedings of the National Academy of Sciences 112, E4854–E4863 (2015).

13. Shao L, Kner P, Rego E H, et al. Super-resolution 3D microscopy of live whole cells using structured illumination. Nature methods 8, 1044–1046 (2011).

14. Shao L, Kner P, Rego E H, et al. Time-lapse two-color 3D imaging of live cells with doubled resolution using structured illumination. Proceedings of the National Academy of Sciences 109, 5311–5315 (2012).

15. Schermelleh, L. et al. Subdiffraction multicolor imaging of the nuclear periphery with 3D structured illumination microscopy. Science 320, 1332–1336 (2008).

16. Smith, C.S. et al. Structured illumination microscopy with noise-controlled image reconstructions. Nature methods 18, 821–828 (2021).

17. Demmerle J. et al. Strategic and practical guidelines for successful structured illumination microscopy. Nature protocols 12, 988–1010 (2017).

18. Zhanghao, K. et al. Super-resolution imaging of fluorescent dipoles via polarized structured illumination microscopy. Nature communications 10, 1–10 (2019).

19. O’Holleran K, Shaw M. Polarization effects on contrast in structured illumination microscopy. Optics Letters 37, 4603–4605 (2012).

20. E. Soubies, M. Unser. Computational Super-Sectioning for Single-Slice Structured-Illumination Microscopy. IEEE Transactions on Computational Imaging 5, 240–250, (2019).

